# Hypergravity is more challenging than microgravity for the human sensorimotor system

**DOI:** 10.1101/2021.12.08.471552

**Authors:** L. Chomienne, P. Sainton, F. Sarlegna, L. Bringoux

## Abstract

While recent findings demonstrated the importance of contextual estimates about gravity for optimal motor control, it remains unclear how gravitational changes are taken into account by the central nervous system to perform complex motor skills. Here, we investigated the effect of microgravity and hypergravity on the neuromuscular control of whole-body reaching movements compared to normogravity. Standing participants (n=9) had to reach toward visual targets during parabolic flights, which allowed us to test the influence of gravity level on sensorimotor planning and control processes. Also, to specifically test the efficiency of online motor control mechanisms, unexpected mechanical perturbations were used. Whole-body kinematics and muscular activity were adjusted in microgravity, allowing arm reaching to be as accurate as in normogravity. In contrast, systematic undershooting was observed in hypergravity, where main parameters of whole-body kinematics remained unchanged and muscle activations insufficiently adjusted to keep the same accuracy as in normogravity. Conversely, muscular synergies exhibited during whole-body reaching were found similar in the various gravitational contexts, as were local muscular adjustments in response to unexpected mechanical perturbations. This suggests that online feedback control remains functional across very distinct gravitational environments. Overall, our findings demonstrates that hypergravity creates challenges that the human sensorimotor system is unable to solve rapidly, contrary to microgravity.

## Introduction

Altered gravity constitutes a problem for the central nervous system because of the fundamental changes of body dynamics which must be integrated to ensure that motor performance is not altered. This integration refers to our capacity to modify the internal representation of the body according to new environmental constraints in order to remain efficient in the given environment. Thereby, understanding how the nervous system is able, or not, to efficiently produce and regulate motor commands in microgravity and hypergravity remains a key issue in the field of human motor control.

As a major feature of our usual surrounding force field, Earth’s gravity has been found to be optimally integrated by the central nervous system (Crevecoeur et al. 2009a; Gaveau et al. 2016, 2021; McIntyre et al. 2001; White et al. 2020; Zago and Lacquaniti 2005). It is well established that the absence of gravitational field results in substantial changes in sensorimotor planning and control (Bock et al. 2003; Lackner and DiZio 2000; Papaxanthis et al. 1998; Tays et al. 2021; Weber et al. 2021). Studying microgravity episodes of parabolic flights has revealed that body unloading is taken into account in the planning of whole-body reaching movements before movement onset (Bringoux et al. 2020; Macaluso et al. 2017). Precisely, the consequences of the novel gravitational context on body dynamics result in a reorganization of motor planning and control, allowing appropriate actions (Berret et al. 2008; Gaveau et al. 2016; Rousseau et al. 2016). Such reorganization, from the earliest stages of exposure to microgravity during parabolic flights, can result in endpoint performance of reaching movements being preserved (i.e., similar as in normogravity; Macaluso et al. 2017; see White et al. 2020 for a review).

Although motor performance has generally found to be unaltered in microgravity, conflicting findings regarding motor responses in hypergravity have been reported. While few studies highlighted unaltered motor performance in hypergravity (e.g., movement accuracy; Crevecoeur et al. 2009; Mierau et al. 2007), some others reported longer or partial sensorimotor reorganization on arm pointing (Bock et al. 1992; Gaveau et al. 2011) and grasping (Crevecoeur et al. 2014) compared to microgravity. For instance, Crevecoeur et al. (2009) reported that, within 3 trials, upward or downward arm movements were correctly executed in hypergravity (i.e., 1.8*g*), as in normogravity. In contrast, Bock et al. (1992) reported a systematic overshot even after 20 parabolas of parabolic flight. Although one might argue that these discrepant observations might be the consequence of the task and underlying mechanisms that could impact motor performance (Crevecoeur et al. 2009a), the overall findings related to human sensorimotor adaptation to hypergravity still appear inconclusive.

Another key issue in altered gravity is how well the central nervous system can deal with unexpected and sudden perturbations. On Earth, the flexibility of sensorimotor control is well known (Franklin and Wolpert 2011; Gomi 2008; Sarlegna and Mutha 2015; Scott 2016; Brenner and Smeets 2018, for reviews). For instance, motor commands can be rapidly adjusted during movement execution in response to a range of mechanical (Cluff and Scott 2013; Crevecoeur and Scott 2013; Kimura and Gomi 2009) or visual (Chomienne et al. 2021; Hayashi et al. 2016; Saijo and Gomi 2010; Telgen et al. 2014) perturbations. In microgravity, Bringoux et al. (2020) reported that an unpredictable target jump triggered at arm movement onset did not affect endpoint accuracy during a whole-body reaching task. This finding highlighted the efficiency of visual feedback control in microgravity. It has been found however that adding a mass on the moving arm could impact endpoint accuracy in both microgravity and hypergravity (Bock et al. 1992). Such type of mechanical perturbation appears to represent a challenge for the central nervous system in altered gravity. However, the online adjustments of muscle activations in response to unexpected mechanical perturbations in novel gravitational contexts remain unclear.

To address these shortcomings, we determined the effect of microgravity and hypergravity on the control mechanisms underlying whole-body reaching movements. To study microgravity and hypergravity, we performed experiments on human participants in an aircraft plane with a specific timing during parabolic flights, resulting in experimentally-controlled changes in background force level, with alternating periods of hypergravity (1.8*g,* nearly twice Earth gravity), normogravity (1*g*), and microgravity (nearly 0*g*). Standing participants were asked to reach with their dominant arm toward visual targets as we recorded endpoint (fingertip) and joint kinematics to determine the multi-joint coordination patterns used in normo-, micro- and hypergravity. We also recorded with surface electrodes the activity of 8 axial and distal muscles to characterize the activation level and synergies in all 3 gravitational contexts. We found that participants precisely reached the targets in microgravity by adjusting muscle activations and kinematics compared to normogravity. In contrast, systematic undershoot was observed in hypergravity, a result of unchanged motor organization relative to normogravity. Also, whatever the gravitational context, final accuracy was maintained to the same level when an unpredictable mechanical perturbation was applied, suggesting that online feedback control processes remain functional in microgravity as well as hypergravity.

## RESULTS

The study was carried out during a 3-day parabolic flight campaign. A programmed sequence of parabolic maneuvers resulted in successive changes of gravitational context that could be decomposed in three phases: 24 s of hypergravity (1.8*g*), 22 s of microgravity (0*g*) and 22 s of hypergravity (1.8*g*; Figure 1D). The normogravity context (1*g*) was present during 1 min between each parabola. In order to study the effect of gravitational context on human motor control, each standing participant was asked at specific times to reach toward one of two visual targets (Figure 1A). These targets were located close (Figure 1B) and far (Figure 1C) to study the planning and control of single-joint shoulder arm movement and whole-body movements (involving the trunk and lower limbs), respectively. In 20% of trials, an electromagnet unexpectedly generated a mechanical perturbation on the wrist at movement onset to specifically study online control mechanisms by means of fingertip and joint kinematics as well as surface electromyography.

**Figure 1:**
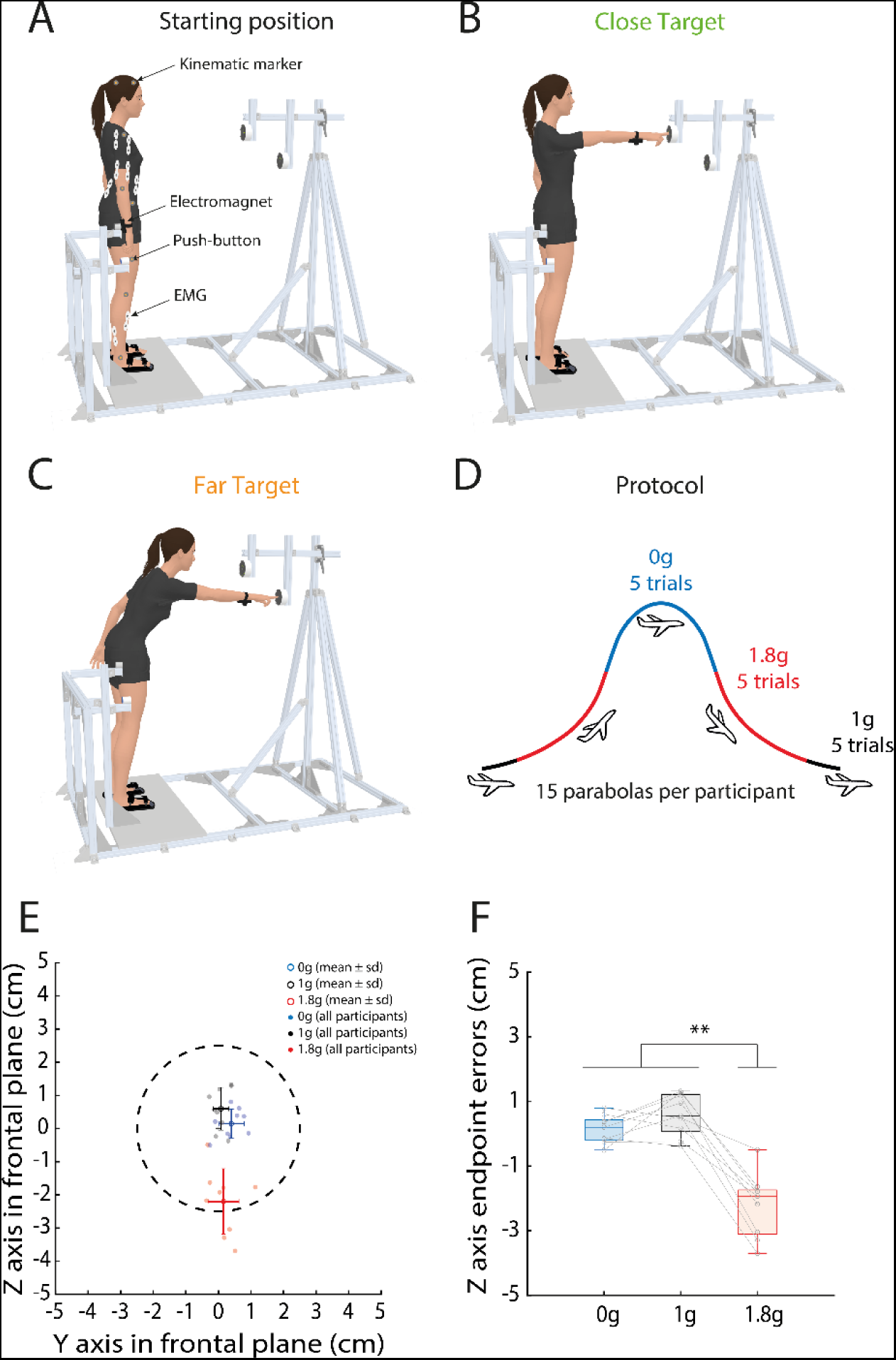
(A) Global view of the experimental set up including the kinematic markers, the EMG electrodes, the push-button to standardize the fingertip start position and the electromagnet to unexpectedly perturb reach movement initiation (20% of trials). Reaching movements were performed toward either a close (B) or far (C) target during parabolic flights which allowed a modification of the gravitational context (D). (E) Endpoint errors as a function of the gravitational environment. The target is represented as a dotted circle (radius: 2 cm). (F) Endpoint errors on Z axis in each gravitational context. Reaching accuracy was impaired in hypergravity (1.8g), as shown by the undershooting relative to the target center. Reaching accuracy was similar in microgravity (0g) and normogravity (1g). **P<0.01.

### Stability of arm reach accuracy across parabolas

How functional is motor performance can be well summarized with final movement accuracy. The accuracy of finger endpoint position relative to the target center was mostly stable across parabolas. Final accuracy was first compared between each parabola (from 1 to 10) in the three different environments (microgravity, hypergravity, normogravity) to determine potential changes across parabolas. A 10x3 ANOVA performed on endpoint error on the Z axis showed a significant main effect of Environment (F_(2, 16)_ = 48.96; p < 0.001; ηp² = 0.86) and a significant interaction between Environment x Parabola (F_(18, 144)_ = 3.09; p < 0.001). Post-hoc analysis revealed that endpoint accuracy in hypergravity (mean = −3.7 cm) only differed in parabola 1 relative to the others (mean = −2.1 cm; p < 0.05) but no significant difference was found between other parabolas, whatever the environment (mean = 4.8 cm). This phenomenon, previously reported in hypergravity (Crevecoeur et al. 2009a), is presumably due to the large initial errors as well as to the rapid adaptation seen in hypergravity. Overall, a systematic undershooting was observed in hypergravity compared to microgravity and normogravity, and the undershooting was significantly greater during the first parabola compared to others. To focus the investigation on the motor responses regarding the gravitational level and avoid initial exposure effects, data from the first parabola were removed (see Sainburg and Kalakanis 2000 for a similar method) and will be specifically discussed elsewhere. In the following sections, we dissect motor performance for the nine last parabolas during which movement accuracy did not significantly differ.

### Endpoint accuracy and arm angular elevation across gravity conditions

Figure 1E shows that final reach errors were greater in hypergravity than in normogravity and microgravity. A 3x2x2 ANOVA [Environment (Hypergravity, Normogravity, Microgravity) x Target (Close, Far) x Perturbation (On, Off)] performed on endpoint errors on the Z axis (perpendicular to the floor) showed a significant main effect of Environment (F_(2, 16)_ = 49.77; p < 0.001; ηp² = 0.86). Figure 1F shows that endpoint errors on the Z axis were greater in hypergravity (-2.2 ± 1.2 cm) than in microgravity (0.1 ± 0.6 cm; p < 0.001) and normogravity (0.6 ± 1.0 cm; p < 0.001). No other main effect or interaction was significant (Target: p = 0.61; Perturbation: p = 0.23; Environment × Target: p = 0.1; Environment x Perturbation: p = 0.98; Target x Perturbation: p = 0.81; Environment x Target x Perturbation: p = 0.85).

The duration of arm movements was influenced by the gravitational environment and the target position. The ANOVA performed on arm movement duration showed a significant main effect of Environment (F_(2, 16)_ = 42.38; p < 0.001; ηp² = 0.84), Target (F(1, 8) = 126.74; p < 0.001; ηp² = 0.94) and a significant interaction between Environment x Target (F(2, 16) = 12.19; p < 0.001; ηp² = 0.6). Obviously, arm movements toward the far target lasted longer (613 ± 90 ms) than those toward the close target (491 ± 60 ms; p < 0.001) whatever the environment. However, movement duration was higher in microgravity (613 ± 105 ms) than normogravity (524 ± 78 ms; p < 0.001) and hypergravity (519 ± 79 ms; p < 0.001), particularly when participants reached the far target (697 ± 76 ms; p < 0.001) as compared to the close target (528 ± 448 ms; p < 0.01). No significant other effects or interactions were found (Perturbation: p = 0.82; Environment × Pertubation: p = 0.99; Target x Perturbation: p = 0.18; Environment x Target x Perturbation: p = 0.80).

The temporal organization of arm reaching was also found to be modified in microgravity. The ANOVA performed on the duration of the acceleration phase, between movement onset and peak velocity, showed a significant main effect of Environment (F_(2, 16)_ = 8.28; p < 0.01; ηp² = 0.51), Target (F_(1, 8)_ = 49.85; p < 0.001; ηp² = 0.86) and Perturbation (F_(1, 8)_ = 5.49; p < 0.05; ηp² = 0.41). Acceleration duration represented a smaller movement duration in microgravity (30 ± 3 %) than in normogravity (3 3± 5 %; p < 0.001) and hypergravity (32 ± 5 %; p < 0.001; Figure 2C). Moreover, acceleration duration was smaller for movements toward the far target (28 ± 4 %; p < 0.001) than the close target (35 ± 4 %; Figure 2G). When the electromagnet was switched on, acceleration duration (31 ± 5 %) was smaller than when it was off (32 ± 5 %; p < 0.05; Figure 2I). No significant interaction was found between these factors (Environment × Target: p = 0.21; Environment x Perturbation: p = 0.79; Target x Perturbation: p = 0.95; Environment x Target x Perturbation: p = 0.79).

**Figure 2:**
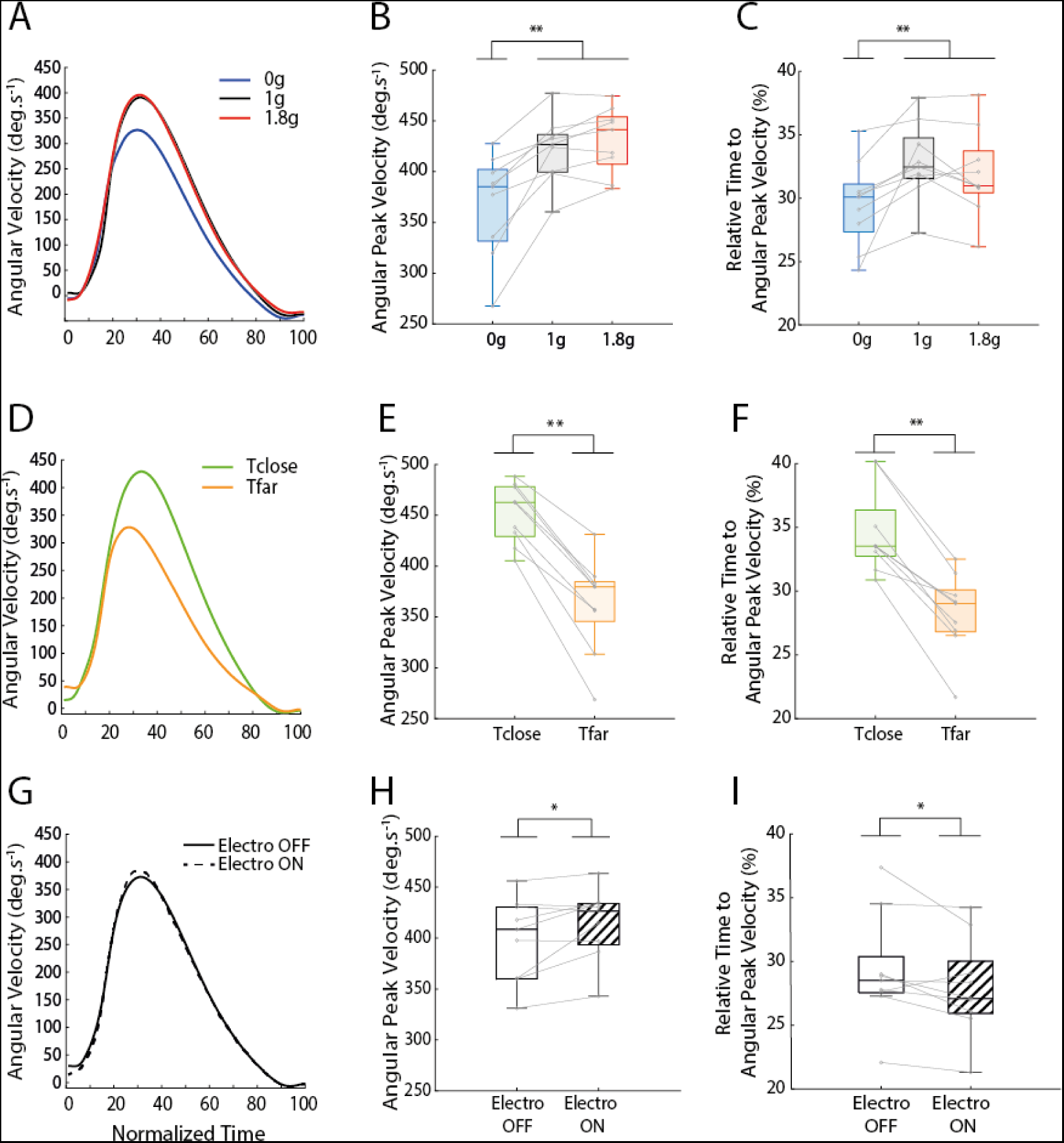
Movement velocity as a function of experimental conditions. (A; D; G) Mean fingertip angular velocity, (B; E; H) Peak Velocity and (C; F; I) relative time to Peak Velocity as a function of (A-C) Environment, (D-F) Target, and (G-I) Perturbation. Environment was either microgravity (0g), hypergravity (1.8g) or normogravity (1g). Participants ether had to reach toward a close or a far target. On 20% of the trials, an electromagnet was switched on and participants had to adjust motor commands to reach the target. *P<0.05, **P<0.01.

Participants modified the spatiotemporal organization of arm movements in microgravity but not in hypergravity with respect to normogravity. The ANOVA performed on peak velocity showed a significant main effect of Environment (F_(2, 16)_ = 30.61; p < 0.001; ηp² = 0.79), Target (F_(1, 8)_ = 81,2; p < 0.001; ηp² = 0.91) and Perturbation (F_(1, 8)_ = 8.3; p < 0.05; ηp² = 0.51). Peak velocity was lower in microgravity (369 ± 73 deg.s^-1^) than in normogravity (424 ± 61 deg.s^-1^; p < 0.001) and hypergravity (433 ± 63 deg.s^-1^; p < 0.001; Figure 2B). Moreover, peak velocity was greater for movements toward the close target (454 ± 48 deg.s^-1^) than the far target (363 ± 61 deg.s^-1^; p < 0.001; Figure 2E). Peak velocity was greater when the mechanical perturbation was unexpectedly switched on (418 ± 71 deg.s^-1^) than when it was off (400 ± 71 deg.s^-1^; p < 0.05; Figure 2H). No significant interaction was found between these factors (Environment × Target: p = 0.26; Environment x Perturbation: p = 0.21; Target x Perturbation: p = 0.61; Environment x Target x Perturbation: p = 0.66).

### Whole-body kinematics

While movements toward the close target were performed as control trials with no requirement to change the body posture to reach the target goal, movements toward the far target were performed to study arm movements and body posture interactions. Indeed, Figure 3A shows that during whole-body reaching, changes in body posture have to be coordinated with the arm movement in order to accurately and rapidly reach a target. Analysis of body orientation was thus performed only for movements toward the far target: it revealed that whole-body reorganizations were present in microgravity but not in hypergravity compared to normogravity. The ANOVA performed on whole-body tilt showed a significant main effect of Environment (F_(2, 16)_ = 27.77; p < 0.001; ηp² = 0.78; Figure 3B). When participants reached toward the far target, whole-body tilt in microgravity (13 ± 3 deg) differed from that in normogravity (9± 2 deg; p < 0.001) and hypergravity (10 ± 2 deg; p < 0.001). No other significant main effect or interaction was found with the other factors (Perturbation: p = 0.31; Environment x Perturbation: p = 0.26). Clearly, a different strategy was used in microgravity compared to normogravity and hypergravity.

**Figure 3:**
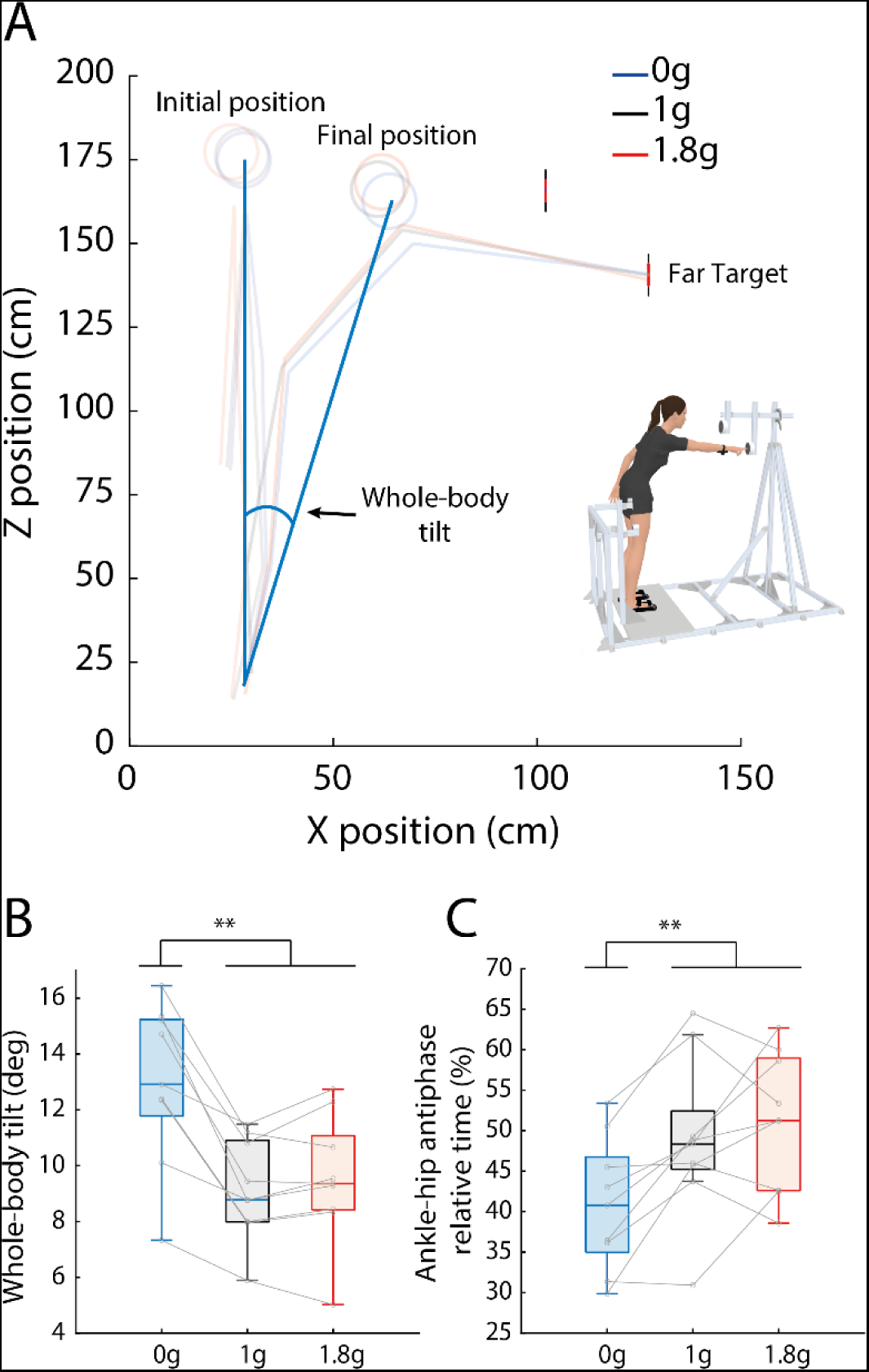
(A) Representative whole-body kinematics during reaching toward the far target for a typical participant in microgravity (0g), hypergravity (1.8g) and normogravity (1g). (B) Mean forward whole-body tilt (orientation of the ankle-head segment relative to vertical) as a function of the Environment. (C) Mean time spent using a hip strategy as a function of the Environment. Whole-body reorganization was only significant in microgravity compared to normogravity. **P<0.01

An ANOVA performed on the time spent by participants using a “hip strategy” (ankle-hip antiphase relative time) showed a significant main effect of Environment (F_(2, 16)_ = 8.61; p < 0.01; ηp² = 0.52; Figure 3C). The relative time using a hip strategy (ankle-hip antiphase relative time) was lower in microgravity (41 ± 80.78 % ± 8.36) compared to normogravity (49 ± 10 %; p < 0.01) and hypergravity (51 ± 9 %; p < 0.01). No other significant main effect or interaction was found with the other factors (Perturbation: p = 0.47; Environment x Perturbation: p = 0.26).

### Muscular synergies

In order to identify synergies between the eight main muscles activated during whole-body reaching, a non-negative matrix factorization was used (NMMF: see Tresch et al. 2006 for more details). The NNMF decomposed the EMG signals in a few synergies (n < M) to describe the contribution of each muscle (from 0 to 1) and the temporal activation pattern of the synergy over time (from 0 to 1). The NNMF computed on the 8 muscles revealed that 3 main synergies were used to control whole-body reaching movements toward the far target (Figure 4). Synergy 1 mainly involved the tibialis anterior and the rectus abdominis around movement onset (Figure 4C). Synergy 2 mainly involved the deltoid anterior and the biceps brachii during arm elevation. Synergy 3 mainly involved the soleus, erector spinae, deltoid posterior, and triceps brachial, mostly around movement offset. To determine whether muscular synergies changed as a function of the gravitational context, a statistical parametric mapping (SPM) was used. The repeated-measure ANOVA conducted with SPM analysis revealed a significant effect of Environment (F_(2, 16)_ = 8.164) only for synergy 1 but post hoc analyses with Bonferroni correction showed no significant difference (p > 0.02).

**Figure 4:**
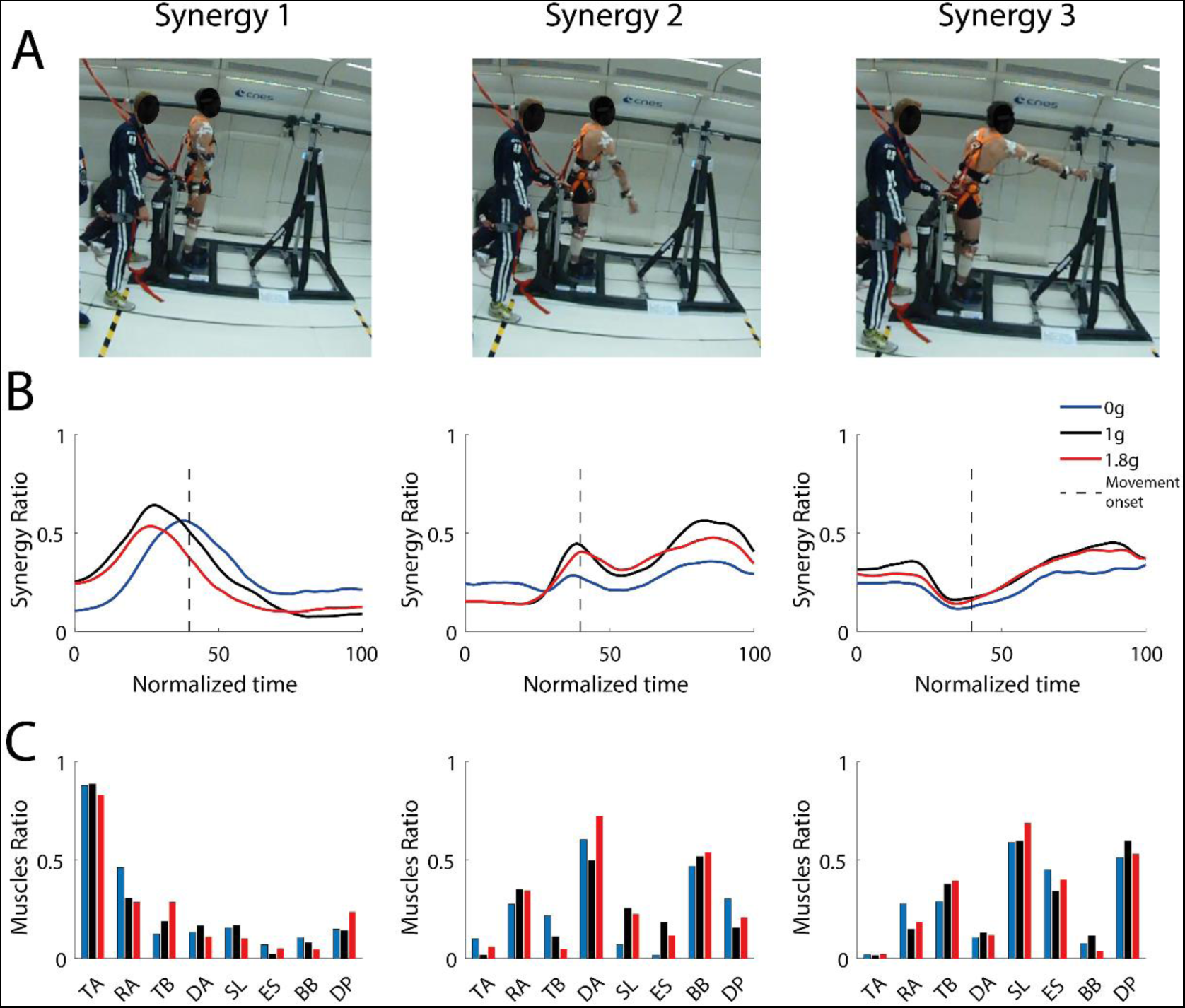
(A) Illustration of three synergies corresponding to three main movement phases in microgravity (0g), normogravity (1g) and hypergravity (1.8g). (B) Mean muscular synergies and (C) muscle ratio for each environment. Synergy 1 corresponded to an activation of Tibial Anterior (TA) and Rectus Abdominis (RA) early in the movement. Synergy 2 in mainly involved the agonist muscles (Deltoid Anterior, Rectus Abdominis, Biceps Brachii) which allowed forward body tilt and arm elevation. Synergy 3 was mainly present at the end of movement where participants slowed down arm elevation by activating antagonist arm muscles (Triceps Brachii and Deltoid Posterior) and maintained final position by activating leg and trunk muscles (Soleus, Rectus Abdominis and Erector Supinae).

### Level of muscle activation

The synergy analysis mainly focused on temporal characteristics of muscular activity. In this second EMG analysis, we aimed at quantifying the level of muscles activation during the planning stage of motor commands to focus on the ability of the nervous system to anticipate the consequences of the gravity environment. We used a 200 ms window from EMG activity onset (see materials and method for more details) and because antagonist muscles are mostly inhibited around movement onset, only the four agonist muscles (tibialis anterior, rectus abdominis, deltoid anterior and biceps brachii) were analysed.

The ANOVA performed on tibialis anterior RMS showed a significant main effect of Environment (F_(2, 16)_ = 8.32; p < 0.01; ηp² = 0.51; Figure 5A). Muscular activity of tibialis anterior was lower in microgravity (196.47 a.u. ± 4.67) than in normogravity (316.25 a.u. ± 76.96; p < 0.01) and hypergravity (271.32 a.u. ± 91.04; p < 0.05). No other significant main effect or interaction was found with the other factors (Perturbation: p = 0.47; Environment x Perturbation: p = 0.23).

**Figure 5:**
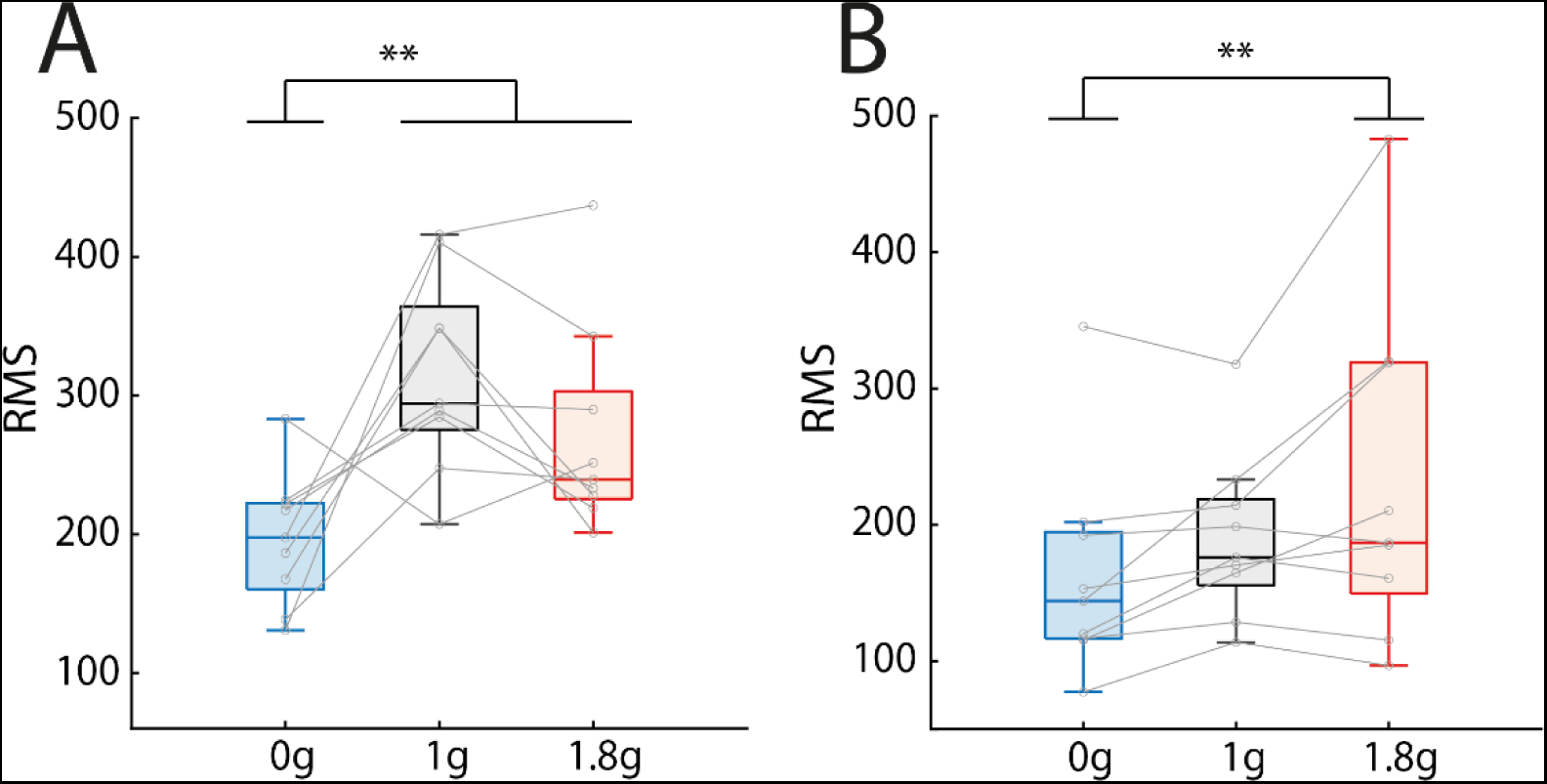
Mean surface EMG root mean square (RMS) of (A) Tibial Anterior (TA) and (B) Biceps Brachii (BB) as a function of Environment. Activity of tibialis anterior was lower in microgravity (0g) than in normogravity (1g) and hypergravity (1.8g) and biceps brachii was less active in microgravity than hypergravity. **P<0.01.

The ANOVA performed on biceps brachii RMS showed a significant main effect of Environment (F_(2, 16)_ = 6.45; p < 0.01; ηp² = 0.45; Figure 5B). Biceps brachii RMS was lower in microgravity (162.92 a.u. ± 93.82) than in hypergravity (230.79 a.u. ± 143.5; p < 0.01). No other significant main effect or interaction was found with the other factors (Perturbation: p = 0.36; Environment x Perturbation: p = 0.31). Furthermore, there was no other significant effect on the other recorded agonist muscles.

## DISCUSSION

The present study determined the effects of microgravity and hypergravity on the motor control mechanisms underlying whole-body reaching. The net output of these mechanisms, as measured by endpoint accuracy, showed little effect of microgravity while a systematic undershoot was found in hypergravity. Analysis of arm and whole-body kinematics as well as muscle activity revealed modifications of motor planning in microgravity compared to normogravity but no significant changes in hypergravity. This supports the view that microgravity can rapidly and efficiently be taken into account by the nervous system to maintain functional motor performance, while hypergravity is a more challenging environment. Despite the fact that online motor control, assessed with responses to unexpected mechanical perturbations, was kept functional in hypergravity, it was not sufficient to minimize reach errors to the level seen in microgravity and normogravity.

### Distinct adapted states in line with the environmental requirements

Microgravity did not affect whole-body reaching performance in terms of final spatial accuracy, as previously reported (Macaluso et al. 2017). This is consistent with previous work (Bringoux et al. 2012; Crevecoeur et al. 2009b; Macaluso et al. 2017; Papaxanthis et al. 2005) suggesting that the optimal integration of such novel body and arm dynamics prior to movement onset allows the central nervous system to produce adapted motor commands in microgravity. This adapted state directly influenced the spatiotemporal organization of arm movement, as, in line with a previous study (Macaluso et al. 2017), peak velocity was observed earlier in the movement. This corresponded to a larger deceleration phase relative to movement duration, which may facilitate sensory feedback control to maintain endpoint accuracy as in normogravity (Sarlegna et al. 2003; Terrier et al. 2011). Consistent with this idea, the present study also revealed a lower peak velocity and a longer movement duration in microgravity, which may reflect participants’ strategy to prioritize accuracy when facing microgravity (Crevecoeur et al. 2010; Mechtcheriakov et al. 2002). In our experiment, the reorganization observed in microgravity could be enhanced by the increased uncertainty caused by successive microgravity and hypergravity episodes during which the task was performed, and the unexpected mechanical constraint which could be present or not at movement initiation.

The analysis of spatial accuracy in hypergravity revealed a systematic undershooting compared to micro and normogravity. This is consistent with previous studies and suggests that, in hypergravity, motor commands are produced according to a normogravity standard and consequently, the force produced to reach the target is too weak (Cohen and Welch 1992; Ross 1991). In the present study, we found similar muscle activations before movement onset in hypergravity and normogravity, and similar joint kinematics and fingertip kinematics. The lack of changes of motor commands in hypergravity compared to normogravity appears clearly related to the observed reach errors, which reveals that hypergravity represents a greater challenge for the human nervous system than microgravity.

Mean endpoint errors appeared at the lower limit of the target (i.e., dotted line; Figure 1E). We speculate that the undershooting observed in the present study as well as in Crevecoeur et al. (2009a) could be explained by participants tending to minimize energy cost while maintaining a reasonable level of performance with respect to the task goals (i.e., “reach the target as fast and as accurate as possible”). A slight change in control policy may be consistent with the optimal control theory (Todorov 2004; Todorov and Jordan 2002), which presents an interesting framework to explain differences in motor performance as a function of the force environment. Whatever the environment, optimality rules usually consist in minimizing energy costs (i.e., by optimizing muscular activations; Berret et al. 2008) while maximizing task success (Harris and Wolpert 1998). However, different environments may be associated with different tradeoffs, hence different control policies and motor performances. The absence of gravitational constraints in microgravity implies a decrease of muscular activity that favors the minimization of energy costs without altering task success. Conversely, the higher gravitational constraint in hypergravity involves higher muscle activations, and minimizing energy costs may involve a reevaluation of task success. By reaching toward the lower limit of the target, participants might have minimized the increase in energy cost typically associated with hypergravity.

### Reorganization of whole-body posture and movement

As previously reported, we found that microgravity resulted in a reorganization of coordination patterns involving the whole body to maintain the performance of reaching arm movements toward the far target. For instance, a larger body displacement was found in microgravity compared to normogravity (Casellato et al. 2012; Macaluso et al. 2017). Moreover, the analysis of hip and ankle joint coordination during movement execution revealed a simultaneous flexion, which corresponds to a preference for an “ankle strategy” (Horak and Nashner 1986) in microgravity which contrasted with the “hip strategy” typically observed in normogravity (Horak 2006; Massion 1992). Because gravitational constraints acting upon the control of stance were absent in microgravity, participants appeared to exploit the possibility of exceeding the usual limits of support surface and showed a larger forward body tilt than in normogravity.

In hypergravity, movement control was similar as in normogravity, across the body. Precisely, analyses of fingertip and joint displacements revealed that arm kinematics, forward trunk displacement and hip and ankle coordination were similar in hypergravity compared to normogravity. While this still allowed participants to maintain balance in hypergravity, we hypothesize that the lack of reorganization at the control level made it difficult to maintain an accurate reach performance, as suggested by the larger reach errors compared to normogravity. Maintaining balance in hypergravity has been reported to involve a greater stiffness of leg joints (Ritzmann et al. 2015) and we speculate that this could be associated with reduced forward body displacement, which may alter manual performance.

### Preserved muscular synergies but subtle adjustments across gravitational contexts

We analyzed muscular synergies to investigate the possible dimensional reduction operated by the central nervous system to produce complex movements in various conditions (d’Avella et al. 2006). Based on an analysis of the activations of the eight muscles mainly involved in the task, three main synergies were found to correspond to three main phases of whole-body reaching movements toward the far target. The first synergy involved an activation of tibialis anterior and rectus abdominis before movement onset. These leg and trunk muscles were engaged to prepare the forward trunk displacement necessary to reach the far target. The second synergy involved an activation of deltoid anterior, rectus abdominis and biceps brachii around movement onset. These agonist muscles were activated to produce both the forward trunk displacement and the arm elevation. The third synergy involved an activation of two groups of muscles toward the end of the movement. The triceps brachial and deltoid posterior were antagonists to the reaching movement, and their activations allowed to slow down the arm elevation and stop at the target. In parallel, leg and trunk antagonist muscles (soleus, rectus abdominis and erector spinea) were activated to maintain final position while the body was tilted forward.

This qualitative analysis of muscular synergies gives a global description of temporal and spatial muscular activity. Interestingly, the number of synergies which best explained all muscular activations remained similar despite the great differences between gravitational contexts. This is consistent with a study of Botzheim et al. (2021) who investigated the effects of gravity on an arm cycling task by comparing two cycling conditions (i.e., sitting and supine). They also found the same number of synergies in both conditions despite considerable biomechanical differences in the gravitational contexts. The similar synergies despite different gravitational contexts could reveal a fundamental building block of whole-body reaching performance. Instead of changing muscular synergies for distinct gravitational constraints, the central nervous system appears to maintain a general structure of muscle activations for movement coordination (Roh et al. 2012). This may minimize the cost of changing the nature of the synergy used most of the time, in normogravity, and appears to work well in microgravity but not as well in hypergravity.

Although synergies were not found to change ‘qualitatively’ across gravitational contexts, we found subtle adjustments of muscle activations as a function of gravity level. Indeed, arm and whole-body kinematics adjustments observed in microgravity compared to normogravity and hypergravity appeared directly linked with context-specific muscular changes. In microgravity, unloading reduces the necessity to produce a high torque at the elbow joint during whole-body reaching movements (Bringoux et al. 2020). Here, because forward body tilt involves lower constraints at the ankle joint in microgravity, we also observed a decrease of tibialis anterior activation at the onset of muscle activation. In hypergravity, only arm kinematic adjustments were found, as revealed by the increased biceps brachii activity around movement onset which allowed to counteract the greater gravitational acceleration. These findings support the idea that hypergravity constraints are somehow taken into account by the nervous system (Crevecoeur et al. 2009a), although insufficiently to perform as in normogravity.

### Preserved online motor control in varied gravitational contexts

Online motor corrections following an unexpected mechanical perturbation were still effective in microgravity and hypergravity to maintain final reach accuracy with respect to movements achieved without perturbation. The perturbation on the wrist around movement onset was associated with an increased velocity of arm elevation. In microgravity, which required participants to deal with the absence of external forces, an unexpected mechanical perturbation was not an obstacle to motor performance. Online motor control was thus efficient, and our findings support and extend those obtained, on a similar task, by Bringoux et al. (2020) who reported efficient feedback responses in microgravity when the target was unexpectedly displaced at movement onset. These findings also support those observed with robotic force fields (Cluff and Scott 2013; Crevecoeur and Scott 2013; Kimura and Gomi 2009) or a modified gravitatoinertial forced field (Chomienne et al. 2021). Overall, the nervous system maintains an impressive ability across various conditions to maintain efficient feedback responses, which is likely linked to the functional interplay between adaptive and fast feedback control mechanisms (Franklin et al. 2012; Gritsenko and Kalaska 2010; Telgen et al. 2014; Wagner and Smith 2008).

## Conclusion

The present study provides new insights on motor performance in altered gravity. Based on different adapted states to gravitational contexts, the central nervous system can maintain motor flexibility in microgravity and hypergravity. However, errors in whole-body reaching as well as the general lack of sensorimotor reorganization observed in hypergravity indicate that hypergravity represents a more challenging state than microgravity for human motor control.

## MATERIALS AND METHODS

### Participants

Nine right-handed volunteers (mean age = 30.8 ± 8.5 years, 4 women) participated in the study. They had no prior experience of parabolic flight and were naïve to the purpose of the study. None of the participants reported any neuromuscular or sensory impairments, as confirmed by a prior medical examination and all had normal or corrected-to-normal vision. Participants gave their signed informed consent prior to the study in accordance with the Helsinki Convention. Before the parabolic flight, participants were given comfort medication (scopolamine) to avoid motion sickness. This medication has been shown to not alter sensorimotor control (e.g., reflex, neuromuscular control, balance performance and force-generating capacity; Ritzmann et al. 2016). The study was authorized by the French National Agency for Biomedical Security (ANSM) and approved by the National Ethic Committee (CPP # 2015-A01231-48).

### Apparatus

Participants stood upright and their feet were fastened to the aircraft floor with foot-straps (Figure 1A). A push-button located alongside their right body at arm length distance from the shoulder was used to standardize the starting fingertip position. In front of each participant, two circular targets (diameter: 4 cm; Figure 1B-C) were located relative to participant’s anthropometry. A close target was set at shoulder’s height at a distance corresponding to the arm length and a far target was positioned 25 cm away and 20 cm below the close target. The positions of the close and far targets were designed to investigate the neuromuscular control of single-joint shoulder arm movement and whole-body movements (i.e., involving movements of the trunk and lower-limb joints), respectively. Participants wore a metal surface bracelet at the wrist, which had to be positioned against an electromagnet located above the push-button (Figure 1A) before movement onset. When activated, the electromagnet could generate a mechanical brake (pullout force: 60 N) at reach initiation. Target illumination and electromagnet activation were controlled with a homemade software (Docometre©) and a real-time acquisition and control system (ADwin-Gold©, Jäger, Lorsch, Germany).

Infra-red active markers were installed on the head, shoulder, hip, knee, ankle, and index fingertip. Their 3D coordinates were recorded at 100 Hz with an optical motion capture system (Codamotion CXS and ActiveHub; Charnwood Dynamics, Leicestershire, UK) for offline analysis of both arm and whole-body displacement.

Surface electromyography (EMG) was recorded at 2000 Hz (BIOPAC Systems, Inc., Santa Barbara, CA) from 4 agonist muscles (Tibial Anterior, TA; Rectus Abdominis, RA; Deltoid Anterior, DA; lateral head of Biceps Brachii, BB) and 4 antagonist muscles (Soleus, SL; Erector Spinae, ES; Deltoid Posterior, DP; short head of Triceps Brachii, TB) involved in the neuromuscular control of whole-body reaching movements. Participants’ skin was cleaned with alcohol and rubbed with an abrasive paper before affixing the surface electrodes (Ag-AgCl; diameter 1 cm, spacing 2 cm) along a line parallel to their fiber orientation to increase the signal-to-noise ratio (according to the SENIAM recommendations; Brindle et al. 2006; Hermens et al. 2000; Mills 2005)

### Procedure

The study was carried out during the #142 parabolic flight campaign of the French national space research center (CNES) including 3 days of flight. The micro- and hypergravity contexts were produced by a series of parabolic maneuvers in a A-310 ZERO-G aircraft chartered by CNES and Novespace. For each participant, a flight involved successive changes of gravitational context that could be decomposed in three phases (Figure 1D): 24 s of hypergravity (1.8*g*), 22 s of microgravity (0*g*) and 22 s of hypergravity (1.8*g*). The normogravity context (1*g*) was present during 1 min between each parabola. Each participant performed 15 trials (5 trials in 0*g*, 5 trials in 1.8*g* and 5 trials in normogravity successively) for each of the 10 parabolas. The total of 150 trials was composed of 75 trials toward the close target and 75 trials toward the far target. In 20% of trials, the electromagnet generated a mechanical brake applied to the wrist while the hand was in start position. Participants were informed of the possible occurrence of these perturbed trials but had no information regarding their sequencing. Targets and Perturbation (i.e., mechanical brake) conditions were presented in a pseudorandom order and counterbalanced between participants.

The task consisted in reaching toward the target with the outstretched arm, as fast and as accurately as possible when the target was switched on. Participants had to maintain the final finger position until target extinction (3 s after movement onset) before returning to the starting position (i.e., upright standing, with the right index finger pressing the start push-button). The experimental session lasted about 20 min.

### Data processing

Data were analyzed using Matlab (Mathworks, Natick, MA). Raw positional data of kinematic markers were low-pass filtered with a dual-pass Butterworth (cut-off frequency: 10Hz; order: 3). Movement duration was defined as the time between movement onset and offset, which were determined when tangential velocity of index finger exceeded and fell below 2% of peak velocity, respectively. The spatial accuracy of reaching movements was analyzed by computing the signed deviation along the longitudinal axis (z-axis) of the right index fingertip at movement offset. Positive and negative longitudinal endpoint error corresponded to overshooting and undershooting, respectively.

To analyze arm kinematics during reaching movement, arm elevation (i.e., shoulder-fingertip angle in the sagittal plane with respect to its initial orientation) was computed across time (Bringoux et al. 2020; Macaluso et al. 2016, 2017). From these values, the angular peak velocity and its relative time of occurrence expressed in percentage of movement duration were computed.

Whole-body movement kinematics and multi-joint coordination patterns were also analyzed. The body displacement at movement offset was computed as the ankle-head angle relative to vertical defined here as perpendicular to the aircraft floor, in the sagittal plane (Figure 3A). The coordination between the ankle and hip joints was examined, using the Continuous Relative Phase analysis (CRP; Bardy et al. 1999; Hamill et al. 2012). For each trial, ankle and hip phase angles were expressed through a “parametric phase plot” where the normalized angular positions (relative to the maximum value of each trial) were plotted relative to the normalized angular velocity. Then, the calculated CRP corresponded to: 𝐶𝑅𝑃(𝑡) = 𝜑_1_(𝑡) − 𝜑_2_(𝑡) where 𝜑_1_(𝑡) and 𝜑_2_(𝑡) are the normalized phase angles for ankle and hip, respectively. Finally, the CRP was scaled from 0° to 180° where a CRP(t) < 90° indicates a preferential ‘ankle strategy’ (ankle and hip moving in-phase) while a CRP(t) > 90° indicates a ‘hip strategy’ (ankle and hip moving in anti-phase). We extracted a single variable from this analysis by computing for each trial the relative time spent by participants using hip strategy (ankle-hip antiphase relative time; expressed in percentage of total movement duration).

Muscular synergies were analyzed, using a non-negative matrix factorization (NNMF; Tresch et al. 2006). Raw EMG data were band-pass filtered with a Butterworth-type filter (cut-off frequency: 20-400 Hz; order: 4) centered around the mean and rectified. Then, a low-pass Butterworth filter (cut-off frequency: 10 Hz; order: 3; Hug 2011) was applied twice (forward and backward to remove phase shift) and the signal was normalized relative to the maximum value of each trial and time was normalized (in % on a time window from −400 ms before movement onset to movement offset). NNMF was computed to identify synergies between the eight muscles. Briefly, the EMG signals were combined into an M × T matrix, where M represents the number of muscles (8 in this study) and T the number of EMG data points (2000) obtained in each trial. The NNMF decomposed the EMG signals in a few synergies (n < M). Each synergy can thus be described as a synergy vector W which represents the contribution level of each muscle (from 0 to 1) and a temporal activation pattern H of the synergy over time (from 0 to 1). The difference between reconstructed and original EMGs was computed using the total variance (tVAF). The maximum iteration of NNMF algorithm was scaled at 1000 times for each number of synergies from 1 to 7 (number of muscles −1), until tVAF increase was < 0.05%. Finally, the number of synergies selected corresponded to a tVAF >90% that is, the number of synergies allows the reconstruction of 90% of the original EMG signal.

In addition, to identify the quantitative changes of muscle activity, root mean square (RMS) of EMG signals was computed for the four agonist muscles (tibialis anterior, rectus abdominis, deltoid anterior, biceps brachial; antagonist muscles are mostly inhibited around movement onset). First, raw EMG data were band-pass filtered with a dual Butterworth (cut-off frequency: 20-400 Hz; order: 4) centered around the mean and rectified. Then, a low-pass Butterworth filter (cut-off frequency: 3 Hz; order: 3) was applied twice (forward and backward to remove phase shift) to create an envelope of the EMG signal (Hug 2011). The activity of each muscle was normalized and expressed as a percentage of the maximum activity observed towards far target trials in normogravity (i.e., baseline condition without disturbance). The onset of EMG activity was identified when the EMG signal exceeded 10% of its peak in the trial. From this onset time, the RMS was computed on a 200 ms window.

### Statistical analyses

To determine the statistical effect of repeated exposure to gravitational changes across the successive parabolas, repeated-measure analyses of variance (ANOVAs) were used to compare the longitudinal endpoint error between each parabola (from 1 to 10) in the different environments (0*g*, 1.8*g*, 1*g*). Subsequently, repeated-measure ANOVAs including 3 environments (0*g*, 1.8*g*, 1*g*) x 2 targets (Tclose, Tfar) x 2 Perturbation conditions (Electro ON, Electro OFF) were performed on arm and whole-body kinematics and RMS EMG variables. Repeated-measure ANOVAs were performed using Statistica software (StatSoft, Inc.). The normal distribution of data for each variable was confirmed by Kolmogorov–Smirnov tests. Posthoc analyses were carried out using Newman–Keuls tests and significance threshold was set at p < 0.05.

To quantify whether muscular synergies changed as a function of the gravitational context a SPM analysis allowing a statistical comparison on the whole time-series of each synergy was used. A repeated-measure ANOVA was conducted on each synergy (1 to 3) for the 3 environments (0*g*, 1.8*g* and 1*g*). SPM analyses were computed in Matlab (MathWorks Inc., Natick, MA) using the open-source software package spm1D (version: M.0.4.7; www.spm1d.org). Significance threshold was set at p < 0.05. Posthoc analyses were performed using SPM paired t-test (0*g* vs. 1.8*g*, 0*g* vs. 1*g*, 1*g* vs. 1.8*g*). Bonferroni correction was used to reduce the statistical risk caused by the multiple tests across these three pairs.

## ACKNOWLEDGMENTS

This work was supported by fundings (DAR 944 & 1013) from the French National Space Research Centre (CNES). The authors thank Sébastien Rouquette (CADMOS) and NOVESPACE for technical support.

## References

d’Avella A, Portone A, Fernandez L, Lacquaniti F. Control of Fast-Reaching Movements by Muscle Synergy Combinations. J Neurosci 26: 7791–7810, 2006.

Bardy BG, Marin L, Stoffregen TA, Bootsma RJ. Postural coordination modes considered as emergent phenomena. J Exp Psychol Hum Percept Perform 25: 1284–1301, 1999.

Berret B, Darlot C, Jean F, Pozzo T, Papaxanthis C, Gauthier JP. The inactivation principle: mathematical solutions minimizing the absolute work and biological implications for the planning of arm movements. PLoS Comput Biol 4: e1000194, 2008.

Bock O, Abeele S, Eversheim U. Sensorimotor performance and computational demand during short-term exposure to microgravity. Aviat Space Environ Med 74: 1256–1262, 2003.

Bock O, Howard IP, Money KE, Arnold KE. Accuracy of aimed arm movements in changed gravity. Aviat Space Environ Med 63: 994–998, 1992.

Botzheim L, Laczko J, Torricelli D, Mravcsik M, Pons JL, Oliveira Barroso F. Effects of gravity and kinematic constraints on muscle synergies in arm cycling. J Neurophysiol 125: 1367–1381, 2021.

Brenner E, Smeets JBJ. Continuously updating one’s predictions underlies successful interception. J Neurophysiol 120: 3257–3274, 2018.

Brindle TJ, Nitz AJ, Uhl TL, Kifer E, Shapiro R. Kinematic and EMG characteristics of simple shoulder movements with proprioception and visual feedback. J Electromyogr Kinesiol 16: 236–249, 2006.

Bringoux L, Blouin J, Coyle T, Ruget H, Mouchnino L. Effect of gravity-like torque on goal-directed arm movements in microgravity. J Neurophysiol 107: 2541–2548, 2012.

Bringoux L, Macaluso T, Sainton P, Chomienne L, Buloup F, Mouchnino L, Simoneau M, Blouin J. Double-Step Paradigm in Microgravity: Preservation of Sensorimotor Flexibility in Altered Gravitational Force Field. Front Physiol 11: 377, 2020.

Casellato C, Tagliabue M, Pedrocchi A, Papaxanthis C, Ferrigno G, Pozzo T. Reaching while standing in microgravity: a new postural solution to oversimplify movement control. Exp Brain Res 216: 203–215, 2012.

Chomienne L, Blouin J, Bringoux L. Online corrective responses following target jump in altered gravitoinertial force field point to nested feedforward and feedback control. J Neurophysiol 125: 154–165, 2021.

Cluff T, Scott SH. Rapid feedback responses correlate with reach adaptation and properties of novel upper limb loads. J Neurosci 33: 15903–15914, 2013.

Cohen MM, Welch RB. Chapter 7 Visual-Motor Control in Altered Gravity. In: Advances in Psychology. Elsevier, p. 153–175.

Crevecoeur F, McIntyre J, Thonnard J-L, Lefèvre P. Movement stability under uncertain internal models of dynamics. J Neurophysiol 104: 1301–1313, 2010.

Crevecoeur F, McIntyre J, Thonnard J-L, Lefèvre P. Gravity-dependent estimates of object mass underlie the generation of motor commands for horizontal limb movements. J Neurophysiol 112: 384–392, 2014.

Crevecoeur F, Scott SH. Priors engaged in long-latency responses to mechanical perturbations suggest a rapid update in state estimation. PLoS Comput Biol 9: e1003177, 2013.

Crevecoeur F, Thonnard J-L, Lefèvre P. Optimal Integration of Gravity in Trajectory Planning of Vertical Pointing Movements. J Neurophysiol 102: 11, 2009a.

Crevecoeur F, Thonnard JL, Lefèvre P. Forward models of inertial loads in weightlessness. Neuroscience 161: 589–598, 2009b.

Franklin DW, Wolpert DM. Computational Mechanisms of Sensorimotor Control. Neuron 72: 425–442, 2011.

Franklin S, Wolpert DM, Franklin DW. Visuomotor feedback gains upregulate during the learning of novel dynamics. J Neurophysiol 108: 467–478, 2012.

Gaveau J, Berret B, Angelaki DE, Papaxanthis C. Direction-dependent arm kinematics reveal optimal integration of gravity cues. eLife 5, 2016.

Gaveau J, Grospretre S, Berret B, Angelaki DE, Papaxanthis C. A cross-species neural integration of gravity for motor optimization. Sci Adv 7: eabf7800, 2021.

Gaveau J, Paizis C, Berret B, Pozzo T, Papaxanthis C. Sensorimotor adaptation of point-to-point arm movements after spaceflight: the role of internal representation of gravity force in trajectory planning. J Neurophysiol 106: 620–629, 2011.

Gomi H. Implicit online corrections of reaching movements. Curr Opin Neurobiol 18: 558–564, 2008.

Gritsenko V, Kalaska JF. Rapid Online Correction Is Selectively Suppressed During Movement With a Visuomotor Transformation. J Neurophysiol 104: 3084–3104, 2010.

Hamill J, Palmer C, Van Emmerik REA. Coordinative variability and overuse injury. Sports Med Arthrosc Rehabil Ther Technol 4: 45, 2012.

Harris CM, Wolpert DM. Signal-dependent noise determines motor planning. Nature 394: 780–784, 1998.

Hayashi T, Yokoi A, Hirashima M, Nozaki D. Visuomotor map determines how visually guided reaching movements are corrected within and across trials. eNeuro 3, 2016.

Hermens HJ, Freriks B, Disselhorst-Klug C, Rau G. Development of recommendations for SEMG sensors and sensor placement procedures. J Electromyogr Kinesiol 10: 361–374, 2000.

Horak FB. Postural orientation and equilibrium: what do we need to know about neural control of balance to prevent falls? Age Ageing 35: ii7–ii11, 2006.

Horak FB, Nashner LM. Central programming of postural movements: adaptation to altered support-surface configurations. J Neurophysiol 55: 1369–1381, 1986.

Kimura T, Gomi H. Temporal development of anticipatory reflex modulation to dynamical interactions during arm movement. J Neurophysiol 102: 2220–2231, 2009.

Lackner JR, DiZio P. Human orientation and movement control in weightless and artificial gravity environments. Exp Brain Res 130: 2–26, 2000.

Macaluso T, Bourdin C, Buloup F, Mille M-L, Sainton P, Sarlegna FR, Taillebot V, Vercher J-L, Weiss P, Bringoux L. Kinematic features of whole-body reaching movements underwater: Neutral buoyancy effects. Neuroscience 327: 125–135, 2016.

Macaluso T, Bourdin C, Buloup F, Mille M-L, Sainton P, Sarlegna FR, Vercher J-L, Bringoux L. Sensorimotor reorganizations of arm kinematics and postural strategy for functional whole-body reaching movements in microgravity. Front Physiol 8: 821, 2017.

Massion J. Movement, posture and equilibrium: interaction and coordination. Prog Neurobiol 38: 35–56, 1992.

McIntyre J, Zago M, Berthoz A, Lacquaniti F. Does the brain model Newton’s laws? Nat Neurosci 4: 693–694, 2001.

Mechtcheriakov S, Berger M, Molokanova E, Holzmueller G, Wirtenberger W, Lechner-Steinleitner S, De Col C, Kozlovskaya I, Gerstenbrand F. Slowing of human arm movements during weightlessness: the role of vision. Eur J Appl Physiol 87: 576–583, 2002.

Mierau A, Girgenrath M, Bock O. Isometric force production during changed-Gz episodes of parabolic flight. Eur J Appl Physiol 102: 313–318, 2007.

Mills KR. The basics of electromyography. J Neurol Neurosurg Psychiatry 76 Suppl 2: ii32–35, 2005.

Papaxanthis C, Pozzo T, McIntyre J. Arm end-point trajectories under normal and micro-gravity environments. Acta Astronaut 43: 153–161, 1998.

Papaxanthis C, Pozzo T, McIntyre J. Kinematic and dynamic processes for the control of pointing movements in humans revealed by short-term exposure to microgravity. Neuroscience 135: 371–383, 2005.

Ritzmann R, Freyler K, Weltin E, Krause A, Gollhofer A. Load Dependency of Postural Control - Kinematic and Neuromuscular Changes in Response to over and under Load Conditions. PLOS ONE 10: e0128400, 2015.

Roh J, Rymer WZ, Beer RF. Robustness of muscle synergies underlying three-dimensional force generation at the hand in healthy humans. J Neurophysiol 107: 2123–2142, 2012.

Ross HE. Motor skills under varied gravitoinertial force in parabolic flight. Acta Astronaut 23: 85–95, 1991.

Rousseau C, Papaxanthis C, Gaveau J, Pozzo T, White O. Initial information prior to movement onset influences kinematics of upward arm pointing movements. J Neurophysiol 116: 1673–1683, 2016.

Saijo N, Gomi H. Multiple motor learning strategies in visuomotor rotation. PloS One 5: e9399, 2010.

Sainburg RL, Kalakanis D. Differences in Control of Limb Dynamics During Dominant and Nondominant Arm Reaching. J Neurophysiol 83: 2661–2675, 2000.

Sarlegna FR, Blouin J, Bresciani J-P, Bourdin C, Vercher J-L, Gauthier GM. Target and hand position information in the online control of goal-directed arm movements. Exp Brain Res 151: 524–535, 2003.

Sarlegna FR, Mutha PK. The influence of visual target information on the online control of movements. Vision Res 110: 144–154, 2015.

Scott SH. A Functional Taxonomy of Bottom-Up Sensory Feedback Processing for Motor Actions. Trends Neurosci 39: 512–526, 2016.

Tays GD, Hupfeld KE, McGregor HR, Salazar AP, De Dios YE, Beltran NE, Reuter-Lorenz PA, Kofman IS, Wood SJ, Bloomberg JJ, Mulavara AP, Seidler RD. The Effects of Long Duration Spaceflight on Sensorimotor Control and Cognition. Front Neural Circuits 15: 723504, 2021.

Telgen S, Parvin D, Diedrichsen J. Mirror reversal and visual rotation are learned and consolidated via separate mechanisms: recalibrating or learning de novo? J Neurosci 34: 13768–13779, 2014.

Terrier R, Forestier N, Berrigan F, Germain-Robitaille M, Lavallière M, Teasdale N. Effect of terminal accuracy requirements on temporal gaze-hand coordination during fast discrete and reciprocal pointings. J Neuroengineering Rehabil 8: 10, 2011.

Todorov E. Optimality principles in sensorimotor control. Nat Neurosci 7: 907–915, 2004.

Todorov E, Jordan MI. Optimal feedback control as a theory of motor coordination. Nat Neurosci 5: 1226–1235, 2002.

Tresch MC, Cheung VCK, d’Avella A. Matrix Factorization Algorithms for the Identification of Muscle Synergies: Evaluation on Simulated and Experimental Data Sets. J Neurophysiol 95: 2199–2212, 2006.

Wagner MJ, Smith MA. Shared internal models for feedforward and feedback control. J Neurosci 28: 10663–10673, 2008.

Weber B, Riecke C, Stulp F. Sensorimotor impairment and haptic support in microgravity. Exp Brain Res 239: 967–981, 2021.

White O, Gaveau J, Bringoux L, Crevecoeur F. The gravitational imprint on sensorimotor planning and control. J Neurophysiol 124: 4–19, 2020.

Zago M, Lacquaniti F. Visual perception and interception of falling objects: a review of evidence for an internal model of gravity. J Neural Eng 2: S198–S208, 2005.

